# SYNGAP1 deficiency disrupts neoteny in human cortical neurons in vivo

**DOI:** 10.1101/2023.01.14.524054

**Authors:** Ben Vermaercke, Ryohei Iwata, Keimpe Weirda, Leïla Boubakar, Paula Rodriguez, Martyna Ditkowska, Vincent Bonin, Pierre Vanderhaeghen

## Abstract

Intellectual deficiency (ID) and autism spectrum disorder (ASD) originate from disrupted development of human-specific cognitive functions. Human brain ontogeny is characterized by a considerably prolonged, neotenic, cortical neuron development. Neuronal neoteny could be disrupted in ID/ASD, but this was never tested because of the difficulties to study developing human cortical circuits. Here we use xenotransplantation of human cortical neurons into the mouse cortex to study the in vivo neuronal consequences of SYNGAP1 haploinsufficiency, a frequent cause of ID/ASD. We find that SYNGAP1 deficient neurons display strong acceleration of morphological and functional synaptic development. At the circuit level, SYNGAP1 haploinsufficient neurons display disrupted neoteny, with faster integration into cortical circuits and acquisition of sensory responsiveness months ahead of time. These data link neuronal neoteny to ID/ASD, with important implications for diagnosis and treatments.

## Introduction

Most neurodevelopmental diseases (NDD) including intellectual disability (ID) and autism spectrum disorders (ASD) lead to deficits in higher cognitive functions, but the underlying mechanisms remain largely unknown (*1*–*4*). Some NDDs could impair species-specific steps of human brain development that underlie the acquisition of higher cognitive functions. One such feature is the human-specific neoteny of cortical neurons, which display considerably prolonged development compared with other mammals (*5*). Cortical neoteny has long been proposed to be important in the acquisition of human-specific neural functions, by extending critical periods of learning and plasticity (*6*). This could be particularly relevant for NDDs that often strike these critical periods (*7*). Synapse and dendritic spine formation and related genes are increased precociously in some ASD patients (*8*, *9*), while early brain overgrowth is a notable feature of many forms of ASD (*10*, *11*).

At least some forms of NDD could thus be causally linked to disruption of human neuronal neoteny, but this has never been tested experimentally given the difficulty to access live human neurons. Indeed the study of NDD mechanisms at the neuronal level in vivo was virtually impossible until recently, because of the lack of a reliable experimental approach to study human neurons integrated into functional circuits in vivo. Recently, new models of xenotransplantation of human cortical neurons in the neonatal rodent cortex have been developed to study human neuron development in vivo (*12*–*15*). Importantly in this system, the transplanted human neurons retain neotenic properties, but are able to synaptically integrate into the mouse cortical circuits and develop functional response properties following their own timeline (*13*).

In the present study we apply xenotransplantation models to study the mechanisms of SYNGAP1 deficiency, one of the most common causes of non-syndromic ID, often associated with ASD and epilepsy (*16*). SYNGAP1 encodes a neuron-specific Ras/Rap-GAP protein that is a key regulator of excitatory synaptic formation and function (*17*), in link with dependent regulation of AMPA receptors (AMPAR) trafficking (*18*–*20*) and dendritic spine morphology (*21*, *22*). Previous work in the mouse has pointed to precocious synapse formation and altered synaptic function and plasticity in the hippocampus and cortex of SYNGAP1 HET mutants (*23*–*26*), making this condition an ideal case to test the potential links between neoteny and ID/ASD in human neurons in vivo.

### Generating an in vivo model of SYNGAP1 deficiency in human cortical neurons

We generated isogenic human pluripotent stem cells (PSC) lines displaying specific heterozygote and homozygote SYNGAP1 mutations using directed genomic engineering. To this aim we inserted mono- or bi-allelic loss of function mutations targeting exon 8, which was chosen because it is common to all *SYNGAP1* alternate transcripts, and is located upstream of the functionally most important functional domains (*17*). Several independent clones displaying mutations on only one allele (heterozygote) or two alleles (homozygote) were selected for phenotypic analysis (Fig. S1A-B). The PSC lines were differentiated into cortical pyramidal neurons in vitro (*12*), which confirmed the loss of SYNGAP1 protein expression of all SYNGAP1 protein isoforms in SYNGAP1 homozygous mutant (hereafter referred to as KO) neurons, and about half reduction in SYNGAP1 haploinsufficient (hereafter referred to as HET) neurons compared with control (hereafter referred to as CTRL) neurons (Fig. S1C). Further characterization of the differentiated neurons in vitro did not reveal detectable changes in neurogenesis or neuronal fate in SYNGAP1 deficient neurons (Fig. 1D and Fig. S1C).

**Fig. 1.**
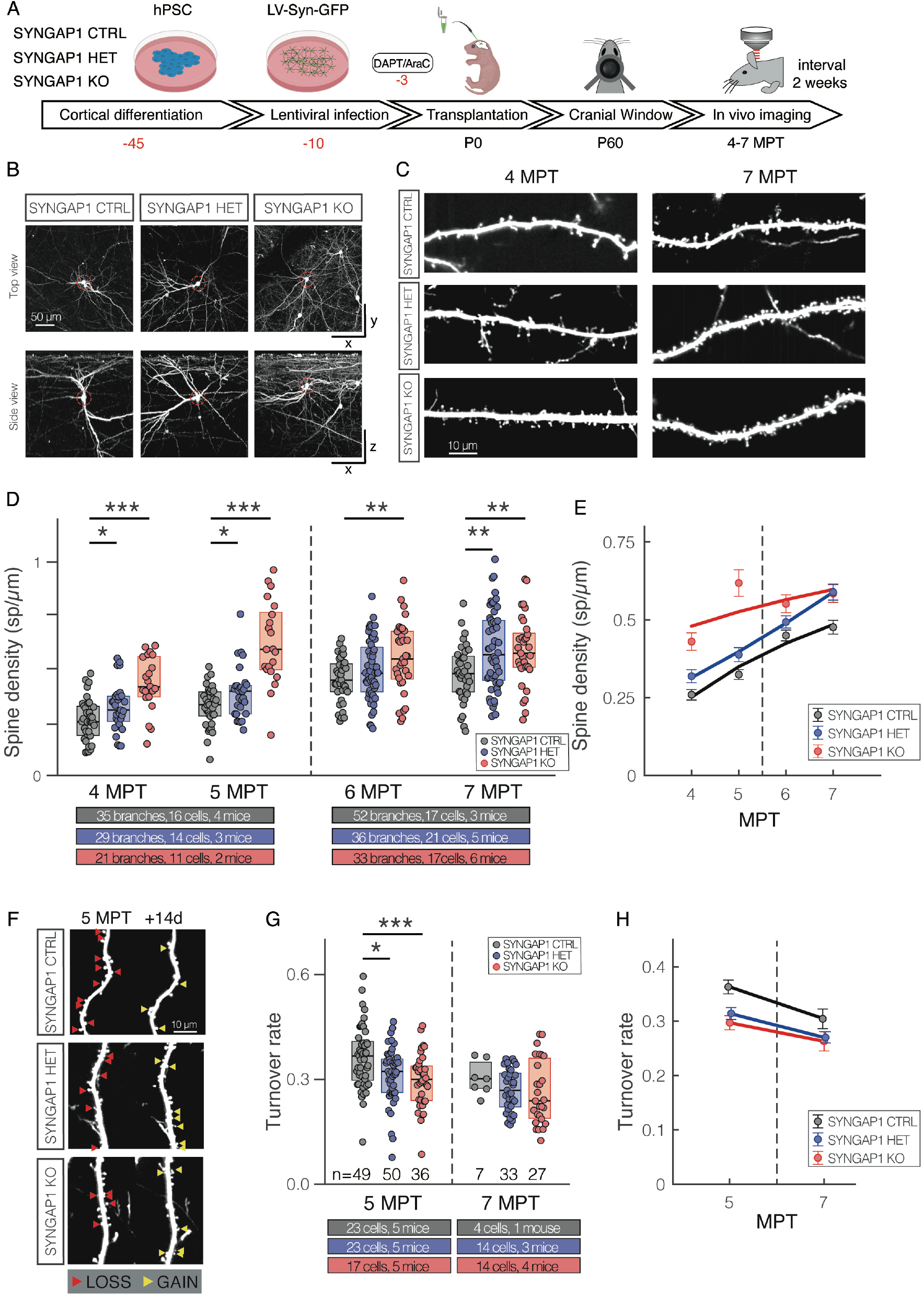
Generating an in vivo model of SYNGAP1 mutation and characterization of spine development and dynamics. **(A)** Experimental timeline: differentiated mutant or control cortical neurons are infected with LV-GFP and transplanted into neonatal mouse pups. A cranial window is implanted in adult mice to allow longitudinal imaging of dendritic branches of transplanted cells. Postnatal day (P), months post-transplantation (MPT). **(B)** Example cell per genotype shown as top and side projection (rows). Total volume size 360×340×400 um. We observe no obvious morphological differences between genotypes. **(C)** Representative dendritic branches per genotype at 4 and 7 MPT. SYNGAP1 KO branches exhibit higher spine density across both timepoints, while SYNGAP HET show a milder phenotype. **(D)** Quantification of spine density for SYNGAP1 CTRL, HET and KO; both mutants differ from controls at early (4 and 5 MPT) and late (7 MPT) timepoints. Data from 4-5 and 6-7 MPT are taken from the same longitudinally sampled branches. Medians per genotype/timepoint: 4MPT: 0.25; 0.31; 0.42, 5MPT: 0.33; 0.40; 0.59, 6MPT: 0.45; 0.48; 0.55, 7MPT: 0.48; 0.57; 0.57. **(E)** Summary of spine density data for all genotypes. The markers and error bars indicate means ± SEMs for each group. The continuous lines are power-law fits (see methods). Note the upward shift of the SYNGAP1 mutants. **(F)** Example of dendritic spine dynamics at 5 MPT for the 3 genotypes. Branches are shown with a 2-week separation. Red arrow heads indicate spines that will disappear at the next timepoint. Yellow arrows indicated spines that are newly formed relative to the previous timepoint. **(G)** Quantification of spine turnover rate (see methods). Both mutants differ from controls at 5 MPT; a similar trend is observed at 7 MPT. Medians per genotype/timepoint: 5MPT: 0.37; 0.32; 0.30, 7MPT: 0.30; 0.27; 0.24. **(H)** Summary of turnover data for all 3 genotypes. Note the downward shift of mutant turnover values compared to controls.

Cortical progenitors were differentiated from PSC of each genotype. Shortly before transplantation they were treated with gamma-secretase inhibitor DAPT and antimitotic agent Ara-C to generate cohorts of neurons born at about the same time prior to transplantation. The neurons were also infected prior to transplantation with ad hoc lentiviral constructs transducing EGFP to enable their identification and morphological analysis. Differentiated human cortical neurons of each genotype were transplanted in newborn (P0) mouse pup littermates, by injecting the cells into the lateral ventricles in the presence of EGTA (*27*). As described previously in this system the newborn neurons then invade the ventricular zone and migrate along the mouse radial glia processes to reach the cortical gray matter (*13*).

### Accelerated dendritic spine formation and stabilization in SYNGAP1 deficient neurons

Next, we examined morphological development of transplanted SYNGAP1 mutant and CTRL neurons with longitudinal in vivo multiphoton imaging of dendritic spines, as a proxy for post-synaptic formation (Figure 1A-C). This enabled us to follow in real time dendritic spine formation and dynamics in transplanted neurons.

Typically, it takes months to develop dendritic spines that stabilize and subsequently reach enough maturity (*13*). Our experiments performed from 4 to 7 months post-transplantation (MPT) revealed accelerated cortical dendritic spine formation in SYNGAP1 deficient neurons. SYNGAP1 KO and HET neurons to a lesser extent displayed higher spine density compared with CTRL neurons, leading to increased values until last time point examined at 7 MPT (Fig 1D,E).

Our longitudinal imaging approach also enable to follow individual spines over weeks and thereby measure the spine turnover ratio, a measure of spine dynamics and plasticity. This revealed similarly decreased dendritic spine turnover ratios in both KO and HET mutant neurons compared with CTRL neurons, indicating that their spines had already reached a high level of stability two months ahead of time (Fig 1F-H). Interestingly, both HET and KO mutant neurons display lower spine turnover rates at 5MPT while KO neurons had much higher spine density (Fig. 1D), suggesting that the observed differences in turnover rates are independent of dendritic spine density.

### Accelerated functional synaptogenesis in SYNGAP1 deficient neurons

We next determined whether the observed increase in the speed of dendritic spine maturation is reflected in neuronal physiological and functional synaptic properties. To this aim *ex vivo* brain slices were prepared from transplanted mice at 4.5 and 6.5 MPT and EGFP-labelled transplanted neurons were examined by patch-clamp recordings. We found no significant differences between the 3 genotypes for all intrinsic properties examined (Fig. 2F-H, Fig.2M-O, Fig. S2), suggesting similar degree of neuronal development. However, we found a robust increase in frequency of excitatory post-synaptic currents (EPSCs) for both mutants, and increased amplitude of EPSCs in HET neurons at early time-points (Fig. 2C,J,K), indicating more numerous and somewhat stronger synapses in SYNGAP1 deficient neurons. We further probed synapse maturity by assessing the ratio of AMPA/NMDA currents, which is known to increase during synaptic development under the control of SYNGAP1 (*17*). We found this ratio to be much increased at 5 MPT in both KO (Fig. 2E) and HET (Fig. 2L), indicative of increased maturation of the post-synaptic compartment. Interestingly the differences in synaptic properties were much more pronounced at early (4.5 MPT) than at later stages (6.5 MPT), consistent with accelerated functional synaptic maturation of the mutant neurons.

**Fig. 2.**
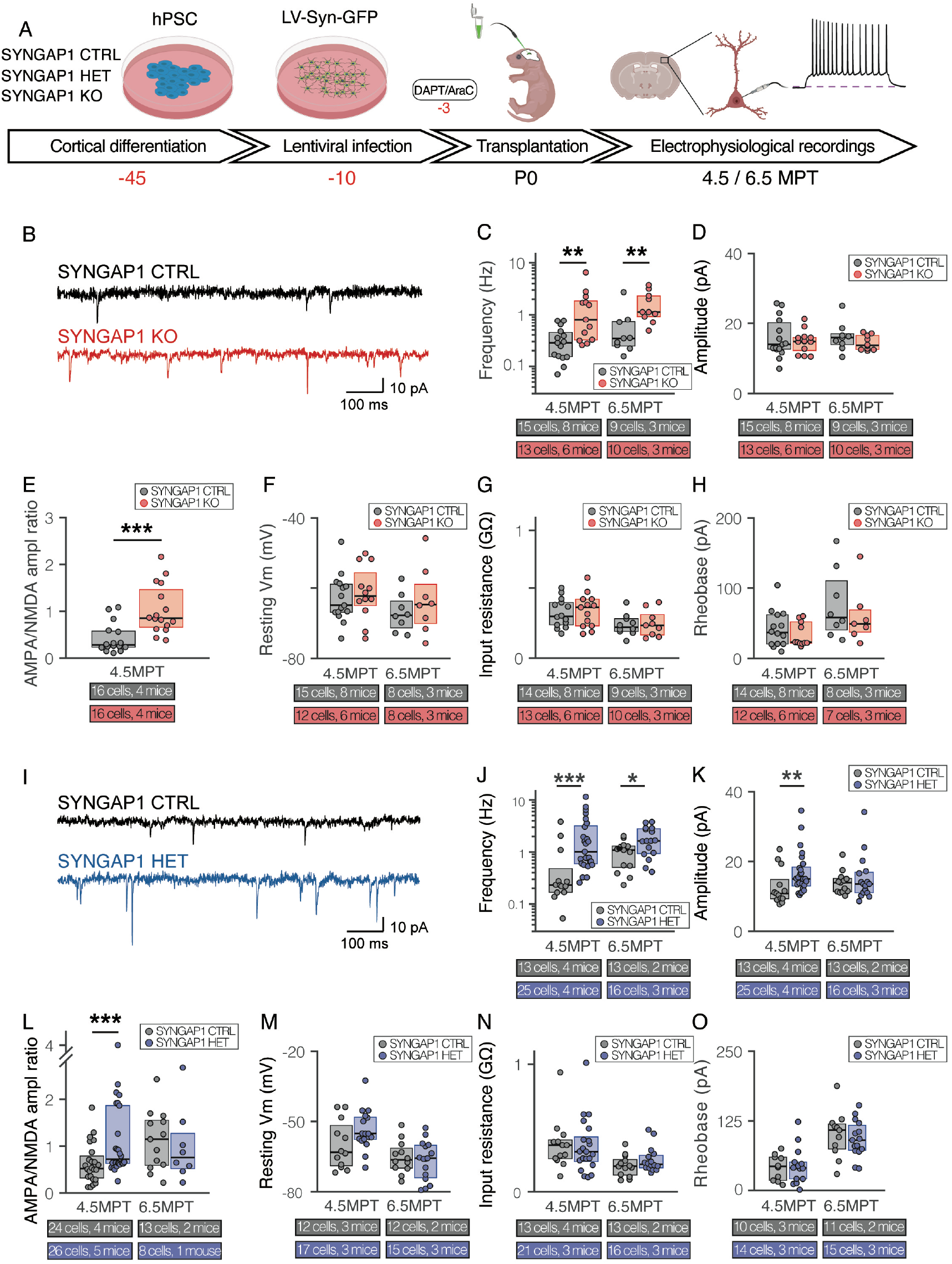
Electrophysiological characterization of SYNGAP1 mutated cortical neurons. **(A)** Experimental timeline: differentiated mutant or control cells are infected with LV-GFP and transplanted. Coronal slices are prepared and GFP labelled cells are targeted for patchclamp experiments. **(B)** Recording traces of SYNGAP1 CTRL and KO neurons showing example mEPSCs. Note more and larger inflections in the red trace. **(C-E)** Quantification of synaptic properties across time: (C) mEPSC frequency at 5 and 7 MPT. Medians per genotype/timepoint: 4.5MPT: 0.28; 0.80, 6.5MPT: 0.35; 1.13. (D) mEPSC amplitude at 5 and 7 MPT. Medians per genotype/timepoint: 4.5MPT: 14.0; 14.80, 6.5MPT: 15.9; 13.7. (E) AMPA/NMDA ratio at 5 MPT. Medians per genotype: 0.28; 0.85. **(F-H)** Cell intrinsic properties show no difference (F) Resting membrane potential, (G) Input resistance and (H) Rheobase. **(I-O)** Same parameters as shown in (A-H) for SYNGAP CTRL and HET at 5 and 7 MPT. (I) Example traces. (J) mEPSC frequency. Medians per genotype/timepoint: 4.5MPT: 0.23; 1.02, 6.5MPT: 1.10; 1.63. and (K) amplitude. Medians per genotype/timepoint: 4.5MPT: 10.8; 15.00, 6.5MPT: 14.0; 13.5. (L) AMPA/NMDA ratio. Medians per genotype/timepoint: 4.5MPT: 0.52; 0.72, 6.5MPT: 1.15; 0.76. (M) Resting membrane potential. (N) Input resistance and (O) Rheobase.

### SYNGAP1 deficient neurons display precocious but normal visual responsiveness

Collectively our data indicate that SYNGAP1 deficient human neurons display accelerated excitatory synaptic development and maturation, suggesting early integration within cortical circuits. To examine consequences at the functional circuit level, we characterized in vivo the development of spontaneous and visually-driven activity of the transplanted neurons in visual cortex (*13*). Neurons in the visual cortex show characteristic sensory response properties such as selectivity for visual orientation and direction. In previous work we showed that about 25% of xenotransplanted neurons develop tuned responses to visual gratings at 6-8 MPT (*13*). We focused on HET neurons for these analyses, given the challenging nature of the experiments and the clinical relevance of haploinsufficiency.

To examine the pace of circuit integration of SYNGAP1 deficient neurons, we therefore probed CTRL and SYNGAP1 HET neurons’ visual response properties by measuring responses to drifting visual gratings (Fig. 3A,B). Longitudinal observations at 2.5-7.5 MPT allowed to detect potential differences in spontaneous and visually-evoked activity.

**Fig. 3.**
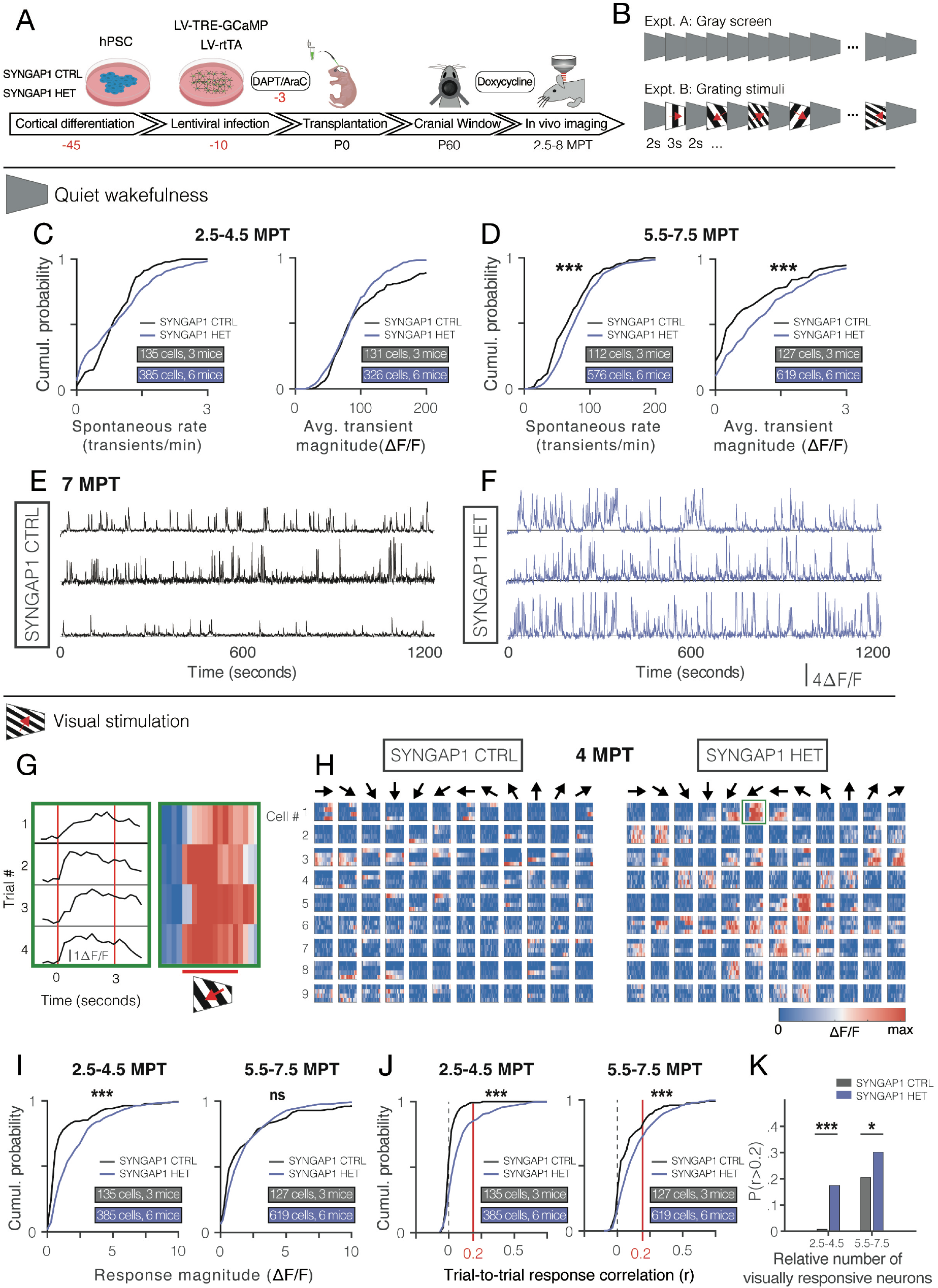
Circuit function in vivo. **(A)** Experimental timeline: differentiated mutant or control cells are infected with LV-TRE-GCaMP/LV-rtTA and transplanted. A cranial window is implanted to allow longitudinal imaging of cellular calcium responses. **(B)** Neurons were stimulated with static gray screen (expt. A) or square-wave drifting gratings of different temporal frequencies, spatial frequencies, spatial orientations, and directions of motion (expt. B). **(C)** Quantification of rate of calcium transients and transient magnitude for SYNGAP1 CTRL and HET neurons recorded between 2.5 and 4.5 MPT. Data from matched FOV between timepoints. Medians per genotype: Rate: 0.88; 0.94, Magnitude: 85.7; 85.2. (D) Transient rate and transient magnitude for control and mutant neurons recorded between 5.5 and 7.5 MPT. Medians per genotype: Rate: 0.40; 0.90, Magnitude: 68.7; 78.0. **(E-F)** Example calcium recordings for 3 cells for SYNGAP1 CTRL (E) and HET (F). Note the increased activity and transient magnitude in the mutants. **(G)** Single trial response to 4 presentations of the same drifting grating pattern. Represented as lines (left) and as a heatmap (right, hot colors are stronger signals). Red solid lines indicate the on- and offset of the stimulus. **(H)** Single trial responses to 4 presentations of all 12 orientations summarized using heatmaps as in (G). The green box marks the example shown in (G). Left shows responses from 9 example SYNGAP CTRL neurons at 4 MPT, right shows responses from 9 example SYNGAP HET neurons. **(I)** Quantification of average response to the preferred stimulus for SYNGAP1 CTRL and HET neurons at both time epochs. Note the large difference in response magnitude at the early time points. Medians per genotype/timepoint: 2.5-4.5: 0.61; 1.61, 5.5-7.5: 0.96; 1.35. **(J)** Quantification of trial-to-trial correlation for SYNGAP1 CTRL and HET neurons at both time epochs. SYNGAP1 mutants show increased values at both time points. Red lines indicate the cut-off value of 0.2 to classify cells as visually responsive. Medians per genotype/timepoint: 2.5-4.5: 0.01; 0.05, 5.5-7.5: 0.03; 0.10. **(K)** Summary of the proportion of visually responsive neurons for SYNGAP1 CTRL and HET at both time points. Proportions per genotype/timepoint: 2.5-4.5: 0.01; 0.18, 5.5-7.5: 0.21; 0.30.

We first examined spontaneous activity in absence of visual stimulation. At 2.5-4.5 MPT, we observed no difference in spontaneous activity in CTRL neurons compared to HET neurons (Fig. 3C). Upon closer inspection, we noted between 2.5 and 3.5 MPT a pronounced correlated spontaneous activity in CTRL neurons, but not in HET neurons (Fig. S3A,B). After 3.5 MPT, this correlated activity disappeared and concomitantly, the activity of CTRL neurons dropped sharply (Fig. S3B bottom). These observations suggest that SYNGAP1 deficient neurons display early desynchronization, while the CTRL neurons show synchronized activity for a longer period, consistent with faster integration of the HET neurons into the mouse cortical circuits (*28*). At later timepoints, HET neurons exhibited more spontaneous calcium transients which were also of greater magnitude (Fig. 3D-F), in line with the increased spontaneous synaptic activity observed using patch-clamp recordings. These data thus indicate that HET neurons display increased integration and activity within the live mouse cortical circuits.

We next analyzed the activity of the transplanted neurons following visual stimulation. More frequent and stronger calcium transients in the HET neurons were observed at both timepoints (Fig. S3C-E). We examined the recorded signals during visual stimulation, focusing the analysis to epochs of visual stimulus (0.5 sec before until 0.5 sec after offset, Fig. 3G), to examine the specificity and sensitivity of the neuron responses. In addition, we correlated the response of each neuron to the direction of moving bars to determine stimulation and/or orientation selective responses (Fig. 3H). This revealed that at early stages 2.5-4.5 MPT HET neurons responded much more strongly than CTRL neurons (Fig. 3I, left panel), while at later timepoints the CTRL neurons increased their responses to the level of HET neurons (Fig. 3I, right panel). These data indicate that the HET neurons display strong acceleration of visual responsiveness at early stages of their development. Increased activity of sensory neurons may be helpful for reliable encoding visual stimuli, but increased baseline activity could also interfere with visual processing. To explore this further we quantified the consistency of the response to four repeated presentations of the same stimulus (Fig. S3F and methods). At both timepoints, we found that SYNGAP1 HET neurons showed higher trial-to-trial correlation (Fig. 3J, Fig. S3G), as well as a higher proportion of visually responsive neurons (Fig. 3K). This suggests that the HET neurons display more robust responsiveness because of stronger synaptic contacts from a potentially more correlated pool of input neurons. These data thus further confirm increased robustness of responsiveness of HET neurons compared with CTRL at early stages of their development, consistent with acceleration of visual functionality. Finally, we found no difference between CTRL and HET neurons for orientation and direction tuning at 7MPT (Fig. S3H,I).

## Discussion

Accelerated timing of synaptic development has been previously associated with some forms of ASD (*8*, *9*, *11*), but whether NDD altering human cognition are actually causally linked to disruption of human neuronal neoteny has remained unclear. Here we have directly tested this hypothesis by focusing on a well-defined ID/ASD form caused by haploinsufficiency of a key synaptic gene frequently mutated in ID/ASD, SYNGAP1, and found strongly disrupted neoteny of human cortical neurons in vivo.

In vitro studies using PSC modeling have previously reported temporal shifts of neurogenesis or early neuronal development with ASD patient-derived cells bearing other gene mutations (*29*, *30*). However in our case, using in vivo modeling of SYNGAP1 alterations, we find a strong and specific acceleration of the developmental tempo of synapses, while earlier development appears to be normal. It remains to be determined whether these differences relate to experimental settings and/or the nature of the genes studied.

Human SYNGAP1 KO PSC-derived induced neurons have been studied in vitro and shown to display enlarged size and increased synaptic activity (*31*). However the consequences of SYNGAP1 pathogenic haploinsufficiency has remained unexplored in human neurons, as well as the consequences of SYNGAP1 KO neurons in vivo, whether in mouse or human. Here we find that that SYNGAP1 KO neurons display some phenotypes to a greater extent than HET mutants, such as increased dendritic spine density, but interestingly dendritic spine turnover and functional synaptic maturation phenotypes are not dramatically different between KO and HET neurons, indicating that the timing of synaptogenesis depends on high levels of SYNGAP1. Importantly our data also strongly point to mostly cell-autonomous effects of SYNGAP1 deficiency, since neotenic disruption happens in isolated human neurons embedded and connected with mouse cortical neurons that contains normal levels of SYNGAP1.

Our data indicate that SYNGAP1 mutant human cortical neurons display a strongly accelerated developmental speed of synaptic development, circuit integration, and sensory responsiveness. The precocious functional integration into cortical circuits of the HET neurons suggest that the cortical circuits in SYNGAP1 affected individuals display a higher rate of synaptic development, thus disrupting the neoteny characteristic of normal human cortical circuits. Interestingly the SYNGPA1 HET human neurons display seemingly normally tuned visual reponses despite their accelerated development, in contrast with observations in mouse SYNGAP1 mutants that were reported to display impaired sensory responses in the barrel cortex (*32*). It will be important to explore in the future the longer term consequences of the disrupted neoteny on final patterning and function of human neurons within neural circuits. These could include the disruption of specific critical periods of plasticity, which are thought to be particularly enhanced and important in our species because of neoteny (*6*). Disrupted neoteny could also lead to miswiring of cortical circuits because of pathologically heterochronic development, as precise timing of neuronal development has a direct impact on patterning of cortical connectivity (*33*, *34*). On the other hand, given the known impact of SYNGAP1 on the function of mature synapses, as well as other phenotypes found in the patients such as epilepsy, it remains to be determined how much mature synaptic defects including an imbalance of excitation/inhibition also contributes to the phenotypes found in the SYNGAP1 patients.

In vitro and in vivo modeling from PSC have emerged as a powerful set of tools to study mechanistically normal and pathological human cortical development. Here we provide a key proof of principle that the mechanisms of NDD can be studied in vivo even at the level of neural circuit function, thus opening novel opportunities to study human neurons integrated in the physiologically relevant context of live cortical circuits.

While the importance of neoteny for human brain development and evolution has long been known (*6*), its relationships with clinical deficits have remained unclear. Our data provide direct in vivo evidence that the disruption of a gene linked to ID/ASD leads to the dysregulation of the tempo of human neuronal development. This could have important implications for the diagnosis and treatments of NDD affected patients.

## Acknowledgments

We thank members of the PV lab and CBD for helpful discussions and precious help. We thank Alexandre Gehanno and Daan Remans for precious help.

## Author contributions

Conceptualization and Methodology, BV, RI, VB, PV; Investigation,.; Formal Analysis,; Writing, BV, PV; Funding acquisition, PV, VB; Resources, PV, VB; Supervision, PV, VB.

## Data and materials availability

All data are available in the manuscript or the supplementary materials. All materials are available upon request from PV or VB.

## Supplementary Materials

### Materials and Methods

#### DNA Constructs

##### pLenti-TRE3G-GCaMP7b-WPRE

The TRE3G fragment (pLenti CMVTRE3G eGFP Puro (w819-1) was a gift from Eric Campeau (Addgene plasmid # 27570; http://n2t.net/addgene:27570; RRID:Addgene_27570)) and the GCaMP7b fragment (pGP-AAV-syn-jGCaMP7b-WPRE was a gift from Douglas Kim & GENIE Project (Addgene plasmid # 104489; http://n2t.net/addgene:104489; RRID:Addgene_104489)) was transferred to lentiviral backbone by restriction enzyme digestion and ligation.

##### pLenti-hSynI-TetON3G-WPRE

The DNA fragment of TetON3G was amplified from AAVS1-TRE3G-EGFP was a gift from Su-Chun Zhang (Addgene plasmid # 52343; http://n2t.net/addgene:52343; RRID:Addgene_52343) using following primers pair: 5’-GTCGACatgtctagactggacaagag-3’ / 5’-ACGCGTttacccggggagcatgtcaa-3’. The TetON3G fragment was transferred to lentiviral backbone pLenti-hSynI (human synapsin I promoter)-WPRE.

##### SYNGAP1 gRNA plasmid

DNA fragment of 5’-TTAGCACATTGTCTACCCGG-3’and 5’-ACGGTACAGATGCAGCCGCA-3’ were transferred to lentiCRISPR v2 plasmid by restriction enzyme digestion and ligation. lentiCRISPR v2 was a gift from Feng Zhang (Addgene plasmid # 52961; http://n2t.net/addgene:52961; RRID:Addgene_52961).

##### SYNGAP1 donor vector (5’HA-EF1α-EGFP-T2A-tCD8-pA-3’HA)

The 5’ homology arm was cloned from H9 hESC DNA using following primers pair: 5’-tGGTACCagcttcctggggctgctata-3’ / 5’-aACGCGTGGCTGTTGTCCTGGCATGG-3’. The 3’ homology arm was cloned from H9 hESC DNA using following primers pair: 5’-tCCTCGAGGaaaGCCGGGTAGACAATGTGCTA-3’ / 5’-CCTAGGTCGCGGAATATGAGGTGCTC-3’. The EF1α promoter-EGFP fragment was prepared from pAAV-EF1α-EGFP/nlsCre-WPRE-pA by restriction enzyme digestion. The T2A-tCD8 fragment was amplified from CD8a-EGFP was a gift from Lei Lu (Addgene plasmid # 86051; http://n2t.net/addgene:86051; RRID:Addgene_86051) using following primers pair: 5’-tTGTACAAGGGATCCGGAGAGGGGAGAGGATCACTGCTGACTTGCGGGGATGTGGAAGAGAACCCAGGGCCCATGGCCTTACCAGTGACC-3’ / 5’-aAGCGCTTCAGTGGTTGCAGTAAAGGGTGATAACCAGTGACAGGAGAAGG-3’. These DNA fragments was transferred to pBlueScript by restriction enzyme digestion and ligation.

#### Human PSC culture and cortical cell differentiation

Human ESC (hESC) (H9; WiCell Cat # NIHhESC-10-0062; female donor) were grown on irradiated mouse embryonic fibroblasts (MEFs) in the ES cell medium until the start of cortical cell differentiation. Cortical cell differentiation was performed as described previously. Two days before starting neuronal cell differentiation, hESCs were dissociated with Accutase (Thermo Fisher Scientific, Cat#00-4555-56) and plate on Matrigel-coated (BD, Cat#354277) plates at low confluency (5,000 cells/cm^2^) in hES medium with 10μM ROCK inhibitor (Merck, Cat#688000). On day 0 of the differentiation, the medium was changed to DDM/B27 medium (DMEM/F12 + GlutaMAX (Thermo Fisher Scientific, Cat#10565042) with N2-supplement (1x, Thermo Fisher Scientific, Cat#A1370701), B27 supplement minus Vitamin A (1x, Thermo Fisher Scientific, Cat#12587010), Bovine Albumin Fraction V (0.05%, Thermo Fisher Scientific, Cat#15260037), 2-Mercaptoethanol (100μM, Merck, Cat#M3148), Non-Essential Amino Acids Solution (1x, Thermo Fisher Scientific, Cat#11140050) and Sodium Pyruvate (1mM, Thermo Fisher Scientific, Cat#11360070)) with recombinant mouse Noggin (100ng/ml, R&D systems, Cat#1967-NG). The medium was changed every other day until day 6. From day 6, the medium was changed every day until day 16. At day 16, medium was changed to DDM/B27 medium and changed every day until day 25. At day 25, the differentiated cortical cells were dissociated using Accutase and cryopreserved in mFreSR (STEMCELL technologies, Cat#05855). Cortical cells were validated for neuronal and cortical markers by immunostaining using antibodies for TUBB3 (1:2,000; BioLegend, Cat#MMS-435P), TBR1 (1:1,000; Abcam, Cat#ab183032), CTIP2 (1:1,000; Abcam, Cat#ab18465), FOXG1 (1:1,000; Takara, Cat#M227), SOX2 (1:2,000; Santa Cruz, Cat#sc-17320), FOXP2 (1:500; Abcam, Cat#ab16046), SATB2 (1:2,000; Abcam, Cat#ab34735), and CUX1 (1:1,000; Santa Cruz, Cat#sc-13024).

#### Establishment of SYNGAP1 mutant cell lines

##### Establishment of SYNGAP1 KO cell lines

hESC were dissociated with Accutase and suspended in Human Stem Cell Nucleofector™ Kit 2 (Lonza, Cat#VPH-5022) with SYNGAP1 gRNA plasmids (5’-TTAGCACATTGTCTACCCGG-3’and 5’-ACGGTACAGATGCAGCCGCA-3’ into lentiCRISPR v2). DNA transfer was performed using Nucleofector II (Lonza) following manufacturer’s instructions. The nucleofected cells were plated on irradiated DR4 MEF (ThermoFisher, Cat#A34966) coated 5cm dishes. At three days after nucleofection, cells were selected with puromycin (500 ng/ml, ThermoFisher, Cat#A1113803) for 72h. At 9-11 days after Puromycin selection, single colony isolation was performed. At 5-7 days after mechanical colony pick-up, expanded cells were cryopreserved in mFreSR and used for DNA extraction. The extracted genomic DNA were used for sequence by MiSeq (Illumina).

##### Establishment of SYNGAP1 HET cell lines

hESC were dissociated with Accutase and suspended in Human Stem Cell Nucleofector™ Kit 2 (Lonza, Cat#VPH-5022) with following mixture: SYNGAP1 donor vector, crRNA/tracrRNA duplex (Integrated Dna Technologies, crRNA sequence: TTAGCACATTGTCTACCCGG) and Alt-R S.p. HiFi Cas9 Nuclease V3 (Integrated Dna Technologies, Cat#1081060). DNA/RNA/protein transfer was performed using Nucleofector II (Lonza) following manufacturer’s instructions. The nucleofected cells were plated on irradiated DR4 MEF coated 5cm dishes. At seven days after nucleofection, cells were used for CD8+ magnetic cell sorting (MACS). The dissociated hESCs were incubated with magnetic beads conjugated anti-human CD8 (Miltenyi Biotec, Cat#130-045-201) in MACS buffer, mixture of autoMACS Rinsing Solution (Miltenyi Biotec, Cat#130-091-222) and MACS BSA Stock Solution (Miltenyi Biotec, Cat#130-091-376), at 4°C for 10 min. CD8 positive selection was carried out with MS column (Miltenyi Biotec, Cat# 130-042-201) according to the manufacturer’s instructions. The sorted cells were plated on irradiated MEF coated plate. At 5-7 days after MACS, expanded cells were cryopreserved in mFreSR and used for DNA extraction. The extracted DNA was used for genomic PCR validation of SYNGAP1 HET/HOMO combined with following primers pair: 5’-TGCAGGACTTTCCAGTTCCC-3’/ 5’-ATCAAGCTGTGGAAGGGTGG-3’.

#### Lentiviral preparation

HEK293T cells were transfected with the packaging plasmids, psPAX2, a gift from Didier Trono (Addgene plasmid # 12260; http://n2t.net/addgene:12260; RRID:Addgene_12260) and pMD2.G, a gift from Didier Trono (Addgene plasmid # 12259; http://n2t.net/addgene:12259; RRID:Addgene_12259), and a plasmid of the gene of interest in the lentiviral backbone: pLenti-hSynI-EmGFP-WPRE; pLenti-hSynI-tdTomato-WPRE; pLenti-UbC promoter-M2rtTA-WPRE (a gift from Rudolf Jaenisch (Addgene plasmid # 20342; http://n2t.net/addgene:20342; RRID:Addgene_20342)); pLenti-hSynI-TetON3G-WPRE (this study); pLenti-TRE-GCaMP6s-P2A-nls-dTomato-WPRE; pLenti-TRE3G-GCaMP7b-WPRE (this study). Three days after transfection, the culture medium was collected and viral particles were enriched by filtering (Amicon Ultra-15 Centrifuge Filters, Merck, Cat#UFC910008). Titer check was performed on HEK293T cell culture for every batch of lentiviral preparation

#### Animals

All mouse experiments were performed with the approval of the KULeuven and ULB Committees for animal welfare. Animals were housed under standard conditions (12 h light:12 h dark cycles) with food and water *ad libitum*. Data for this study are derived from a total of 65 mice of both sexes.

#### Xenotransplantation

Human neuron xenotransplantation was performed as described previously. Human cortical cells (frozen at day 25) were thawed and plated on Matrigel-coated plates using DDM/B27+Nb/B27 medium at 37°C with 5% CO_2_. At seven days after thawing (day 32), cells were dissociated using Accutase and plated on Matrigel-coated plate at high confluency (450,000-700,000 cells/cm^2^) with lentiviral vector: LV-hSynI-EmGFP-WPRE or LV-hSynI-tdTomato-WPRE for spine imaging; LV-UbC-M2rtTA-WPRE / LV-TRE-GCaMP6s-P2A-nls-dTomato-WPRE or LV-hSynI-TetON3G-WPRE / LV-TRE3G-GCaMP7b-WPRE for functional imaging. Nine days later (day 41), the medium was changed to DDM/B27+Nb/B27 medium with 10μM DAPT (Abcam, Cat#ab120633) and cultured for two additional days. At day 43, the cells were treated with 5μM Cytarabine (Merck, Cat#C3350000) for 24h. On the following day, the cortical cells were dissociated using NeuroCult™ Enzymatic Dissociation Kit (STEMCELL technologies, Cat#05715) following manufacturer’s instructions and suspended in the injection solution containing 20mM EGTA (Merck, Cat#03777) and 0.1% Fast Green (Merck, Cat#210-M) in PBS at 100,000–200,000 cells/μl. Approximately 1-2μl of cell suspension was injected into the lateral ventricles of each hemisphere of neonatal (postnatal day 0 or 1) immunodeficient mice (Rag2^-/-^) using glass capillaries pulled on a horizontal puller (Sutter P-97).

#### Western blot

Human cortical cells were collected and lysed in RIPA buffer (Cell Signaling Technology, Cat#9806) with 1x Protease inhibitor (Roche, Cat#11873580001) at 4°C with rotation. Protein concentration was measured using Bradford assay (Thermo Fisher). The sample were run in a NuPAGE^®^ 4-12% Bis-Tris Gel (Thermo Fisher, Cat#NP0321) at 90 V for 2h and then transferred to Immun-Blot^®^ PVDF Membrane (BioRad) at 100 V for 2h. The membrane was blocked in TBS supplemented with 5% skim milk and 0.1% TWEEN^®^ 20 (Sigma-Aldrich) for 1h at room temperature and subsequently incubated overnight at 4°C with primary antibodies (SYNGAP1 (1:1,000, Thermo Fisher, Cat#PA1-046) and GAPDH (1:5,000, Sigma-Aldrich, Cat#G8795)) diluted in the blocking solution. The membrane was washed three times and incubated with adequate HRP-conjugated secondary antibody for 1h at room temperature. The signal was detected by Western Lightning Plus-ECL, Enhanced Chemiluminescence Substrate (PerkinElmer).

#### Electrophysiological recordings

Whole cell patch-clamp recordings were performed on acute coronal slices prepared from 6 months old mice with xenotransplanted PSC-derived human neurons. Briefly, animals were anaesthetized intraperitoneally with Nembutal and transcardially perfused with ~25 mL of icecold NMDG-based slicing solution. Brains were rapidly extracted and placed in ice-cold NMDG-based slicing solution containing (in mM): 93 N-Methyl-D-glucamine, 2.5 KCl, 1.2 NaH2PO4, 0.5 CaCl2, 10 MgSO4, 30 NaHCO3, 5 Na-ascorbate, 3 Na-pyruvate, 2 Thiourea, 20 HEPES and 25 D-glucose (pH adjusted to 7.35 with 10 N HCl, gassed with 95% O2/5% CO2). Coronal slices (250 μm) were cut in ice-cold NMDG-based slicing solution (using a Leica VT1200) and subsequently incubated for ~6 minutes in the NMDG solution at 34°C. Slices were then transferred into holding aCSF, containing (in mM): 126 NaCl, 3 KCl, 1 NaH2PO4, 1 CaCl2, 6 MgSO4, 26 NaHCO3 and 10 D-glucose (gassed with 95% O_2_/5% CO_2_). Slices were stored at room temperature for ~1 hour before experiments.

During experiments brain slices were continuously perfused in a submerged chamber (Warner Instruments) at a rate of 3-4 ml/min with 127 mM NaCl, 2.5 mM KCl, 1.25 mM NaH 2 PO 4, 25 mM NaHCO 3, 1 mM MgCl 2, 2 mM CaCl 2, 25 mM glucose at pH 7.4 with 5% CO 2 / 95% O 2. For sEPSC and AMPA/NMDA ratio recordings we added 20 μM bicuculline. Whole cell patch clamp recordings were done using borosilicate glass recording pipettes (resistance 3.5–5 MΩ, Sutter P-1000) filled with the following internal solution: 115 mM CsMSF, 20 mM CsCl, 10 mM HEPES, 2.5 mM MgCl 2, 4 mM ATP, 0.4 mM GTP, 10 mM Creatine Phosphate and 0.6 mM EGTA. Visually identifiable fluorescently labelled transplanted neurons were selected for recording. Whole-cell patch-clamp recordings were done using a double EPC-10 amplifier under control of Patchmaster v2 x 32 software (HEKA Elektronik, Lambrecht/Pfalz, Germany). Currents were recorded at 20 Hz and low-pass filtered at 3 kHz when stored. The series resistance was compensated to 75-85%. Spontaneous input was recorded using whole-cell voltage clamp recordings (Vm =-70 mV). An extracellular stimulation pipette (borosilicate theta glass, Hilgenberg) was placed near the recorded neuron and used for initiating evoked AMPAR and NMDAR-mediated currents (80-120 μA, 1ms (Isoflex, A.M.P. Instruments LTD)). AMPAR-mediated evoked EPSCs were measured in whole-cell voltage clamp at a holding potential of −70 mV, while the NMDAR-mediated component was measured at +40 mV immediately after the initial AMPAR/NMDAR-mediated current (100-150 ms after electrical stimulation). Evoked data were analysed using Fitmaster (HEKA Elektronik, Lambrecht/Pfalz, Germany), spontaneous input was analyzed using Mini Analysis program (Synaptosoft).

#### Surgical procedures

Standard craniotomy surgeries were performed to gain optical access to the visual cortex through a set of cover glasses (*34*). Rag2KO mice aging between 2 and 6 months were anaesthetised (isoflurane 2.5%–3% induction, 1%–1.25% or a mix of ketamine and xylazine 100mg/kg and 10 mg/kg respectively). A custom-made titanium head plate was mounted to the skull, and a craniotomy over the visual cortex was made for calcium imaging. The cranial window was covered by a 5 mm cover glass. Buprenex and Cefazolin were administered postoperatively (2 mg/kg and 5 mg/kg respectively) when the animal recovered from anaesthesia after surgery.

#### Widefield calcium imaging

Widefield fluorescent images were acquired through a 2x objective (NA = 0.055, Edmund Optics). Illumination was from a blue LED (479nm, ThorLabs), the green fluorescence was collected with an EMCCD camera (EM-C2, QImaging) via a bandpass filter (510/84 nm filter, Semrock). The image acquisition was controlled with customized software written in Python.

#### Two-photon calcium imaging

A customized two-photon microscopy (Neurolabware) was used. GCaMP6s were excited at 920 nm wavelength with a Ti:Sapphire excitation laser (MaiTai eHP DeepSee, Spectra-Physics). The emitted photons were split by a dichroic beamsplitter (centered at 562 nm) and collected with a photomultiplier tube (PMT, Hamamatsu) through a bandpass filter (510±42 nm, Semrock) for the green fluorescence of GFP or GCaMP6s and a bandpass filter (607±35 nm, Semrock) for the red fluorescence of nls-dTomato.

#### In vivo structural imaging

Animals were aneasthetised using a mix of ketamine (100 mg/kg) and xylazine (10mg/kg) at 1% ml/g of their body weight. They were placed on a sterilized plexiglass platform and protective eye ointment was applied to both eyes.

The high NA objective (25x Olympus, 1.05 NA) was aligned to be orthogonal to the surface of the top coverslip. All imaging was performed by moving the motorised stages of the microscope along a virtual axis parallel to the optical axis of the objective. For each neuron, we collected a 3D overview stack spanning 360 x 340 x 450 μm. We then repeatedly acquired high resolution anatomical stacks from anaesthetised mice. Typical stacks consisted of 200-300 optical section spaced 1 μm apart. The imaged area spanned 127 x 110 μm. We recorded both green and red channels which allows us to separate GFP signals from endogenous fluorescence. To reduce effects of shot noise, we averaged 50 frames collected per section. Using these preprocessed stacks, we traced branches of interest using a custom matlab GUI. We then extracted 3D sub-volumes (28 x 20 x X μm, where X is the length of the branch, ranging 50-120 μm) rotated in all 3 dimensions to closely fit each branch. These sub-volumes were used for subsequent analysis.

#### In vivo functional imaging

To enable functional imaging of transplanted human cells, these were infected *in vitro* with both LV-TRE-GCaMP6s-P2A-nls-dTomato-WPRE and LV-UbC-M2rtTA-WPRE or LV-TRE-GCaMP7s and LV-UbC-M2rtTA-WPRE two weeks ahead of transplantation. In these conditions, 50%–70% cells express GCaMP (and nls-dTomato) under doxycycline treatment. We started doxycycline treatment as of two weeks before the imaging experiments.

Two-photon images (702×796 pixels per frame) were collected at 20 Hz with a 16x objective (Nikon 0.80 NA). Volume imaging was accomplished by using an electro-tunable lens (EL-10-30-TC, Optotune) to move the focal plane using a sawtooth pattern in 4 or 5 steps (50 μm separation). We simultaneously recorded neuronal activities in large volumes (1 × 1.5 × 0.20 mm^3^) of layer 2/3 visual cortex. During imaging, mice were head-clamped on a sterile plexiglass platform while consciously viewing the visual stimuli on the display. Eye movements were monitored using a camera and infrared illumination (720–900 nm bandpass filters, Edmund).

Visual stimuli were displayed on a gamma-corrected LCD display (22’’, Samsung 2233RZ). The screen was oriented parallel to the eye and placed 18 cm from the animal (covering 80 degrees in elevation by 105 degrees in azimuth). Spherical correction was applied to the stimuli to define eccentricity in spherical coordinates. Visual stimuli consisted of drifting square wave gratings in 6 combinations of 3 spatial frequencies (SF=0.04, 0.08, 0.16 cycles per degree) and 2 temporal frequencies (TF=1, 4 Hz) in 12 directions covering 360 degrees. All 72 stimuli were presented in random order and this block of 72 stimuli was repeated 4 times.

## QUANTIFICATION AND STATISTICAL ANALYSIS

### Electrophysiology

All electrophysiological recordings were analyzed using MATLAB (The Mathworks, Natick, MA). Raw voltage traces were filtered at 5 kHz and passive membrane and AP properties were extracted and saved in a database for further analysis and statistics.

### In vivo spine imaging

Segmented branches for adjacent time points were loaded into a custom GUI and all protrusions were marked as spines. During this process we took the 3d structure of the spine into account by scrolling through the volume. This allowed us to distinguish between actual spines and processing passing above and below the branch of interest. We then correlated spines in both time points using their location relative to branch landmarks. Spines that were present in both time points were marked as *conserved*. Spine present only in the first time point were marked as *lost* and, conversely, spines that were found only in the second time point were marked *gained*. Spines present on a non-overlapping section of the branch were marked, but only included for the spine density quantification. The spine *density* was calculated by counting the number of actual spines on each segment and dividing this count by the length of the branch. This length was obtained by summing the distances between adjacent markers used to trace the branch. Spine *turn-over* was calculated by dividing the sum of gained and lost spines by the sum of spines present at both time points. The resulting time lapse curves were analysed using a mixed model ANOVA with age group as between-subject factor and time point as within-subject factor. Spine density for gained and lost spines was calculated by counting gained and lost spines and dividing by branch length.

### Calcium imaging

Two-photon movies for all experiments collected during 1 session were motion registered to a common reference image. This image was constructed by registering and then averaging 1200 frames from the middle of the session. We manually segmented regions-of-interest (ROIs) using information from both red and green channels. We then extracted cellular time courses for each ROI by averaging all pixels inside each ROI, removing the neuropil signal and correcting for slow baseline drift. We then calculated ΔF/F_o_ traces by subtracting the baseline fluorescence (F_o_) corrected time course and dividing by F_o_.

To quantify activity of neurons, we used a differential algorithm that detected epochs were the calcium trace increased by more than 2x STD/second, where STD is the standard deviation of the distribution constructed by combining the negative part of the trace with its sign-inverted counterpart. Neurons were considered *active* if their calcium trace exhibited more than 0.5 transients per minute. Neurons were deemed *visually responsive* if, the trial-to-trial response correlation (see Fig. S3F) exceeded 0.20.

For orientation tuning experiment we determined the *orientation selectivity index* (OSI) and *direction selectivity index* (DSI) based on circular variance as proposed by (*35*).

### Statistical analysis

Results are shown as mean±standard error (S.E.M.) of at least three biologically independent experiments. Electrophysiological properties are shown as median and interquartile range, as indicated in figure legends. Student’s unpaired *t*-test was used for two group comparisons. Analyses of multiple groups were performed by a one-way or two-way analysis of variance (ANOVA) followed by post hoc multiple comparisons using Tukey’s test. Additional statistical details are given in the figure legends and in the main text. The number of samples N can be found in the figures.

---

**Fig. S1.**
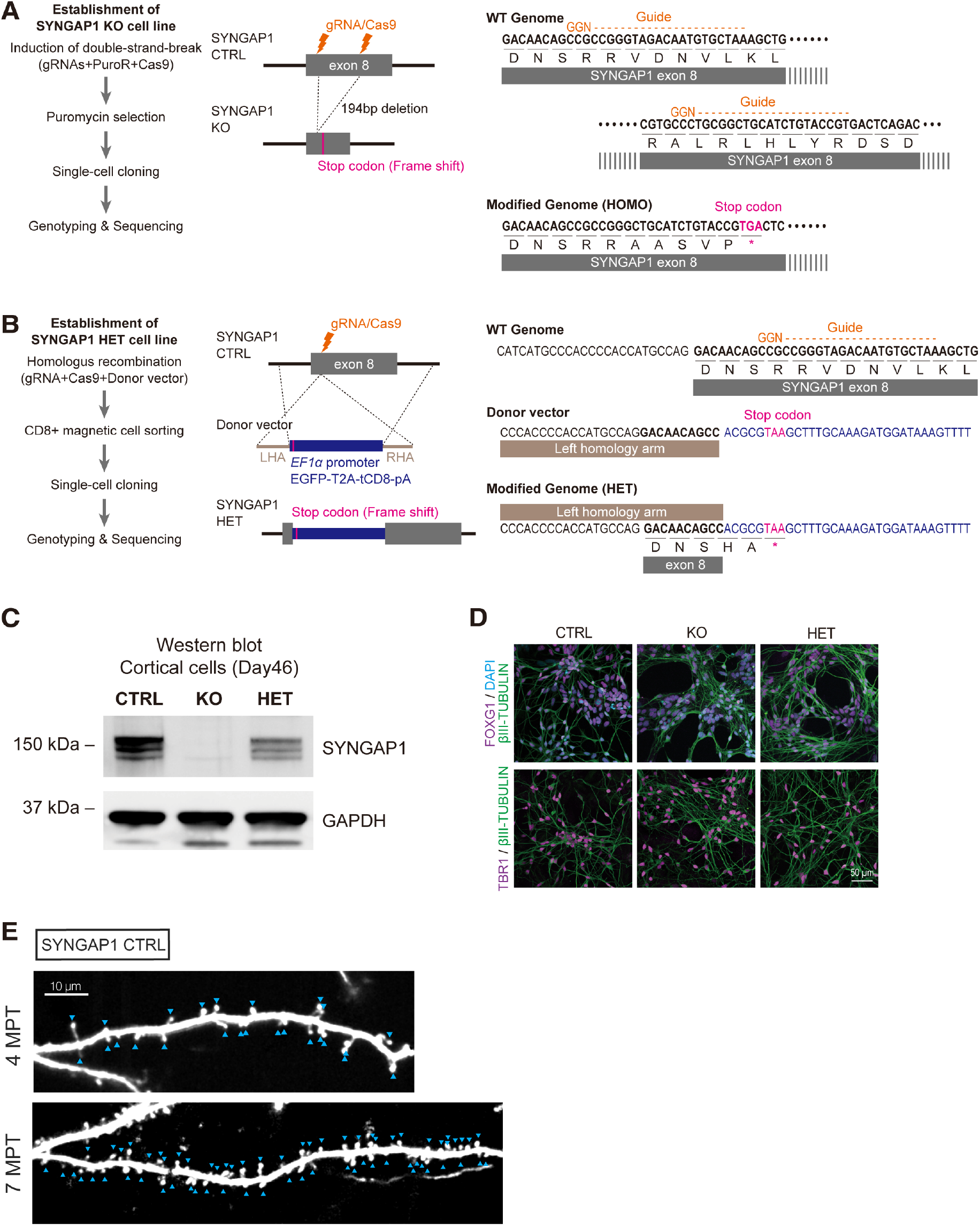
Establishment of SYNGAP1 mutant cell lines. **(A)** The schema of SYNGAP1 KO cell line establishment. Two gRNAs, Cas9 gene and Puromycin resistance (PuroR) gene expressing plasmids were transfected by nucleofection into human PSCs. The cells were selected by Puromycin treatment and used for single-cell cloning. The 194-bp deletion in exon8 induced a frameshift. Right: target sites of gRNAs. **(B)** The schema of SYNGAP1 HET cell line establishment. Single gRNA, Cas9 protein and donor vector were transfected by nucleofection into human PSCs. The cells were used for CD8 positive magnetic sorting. The donor vector sequence in exon 8 induced a frameshift. Right: target site of gRNA. **(C)** Representative image of Western blot for SYNGAP1 protein from whole cell protein extracts of human PSC-derived cortical cells at day 46. **(D)** Representative images of cortical marker positive cortical cells at day 32 after differentiation. (Top) Telencephalon-specific marker FOXG1. (Bottom) Deep-layer marker TBR1. Scale bar: 50 μm. **(E)** Example of dendritic spine quantification. Two branches taken from SYNGAP1 CTRL neurons at 4 and 7 MPT are shown with blue arrow heads indicating structures that we identified as dendritic spines.

**Fig. S2.**
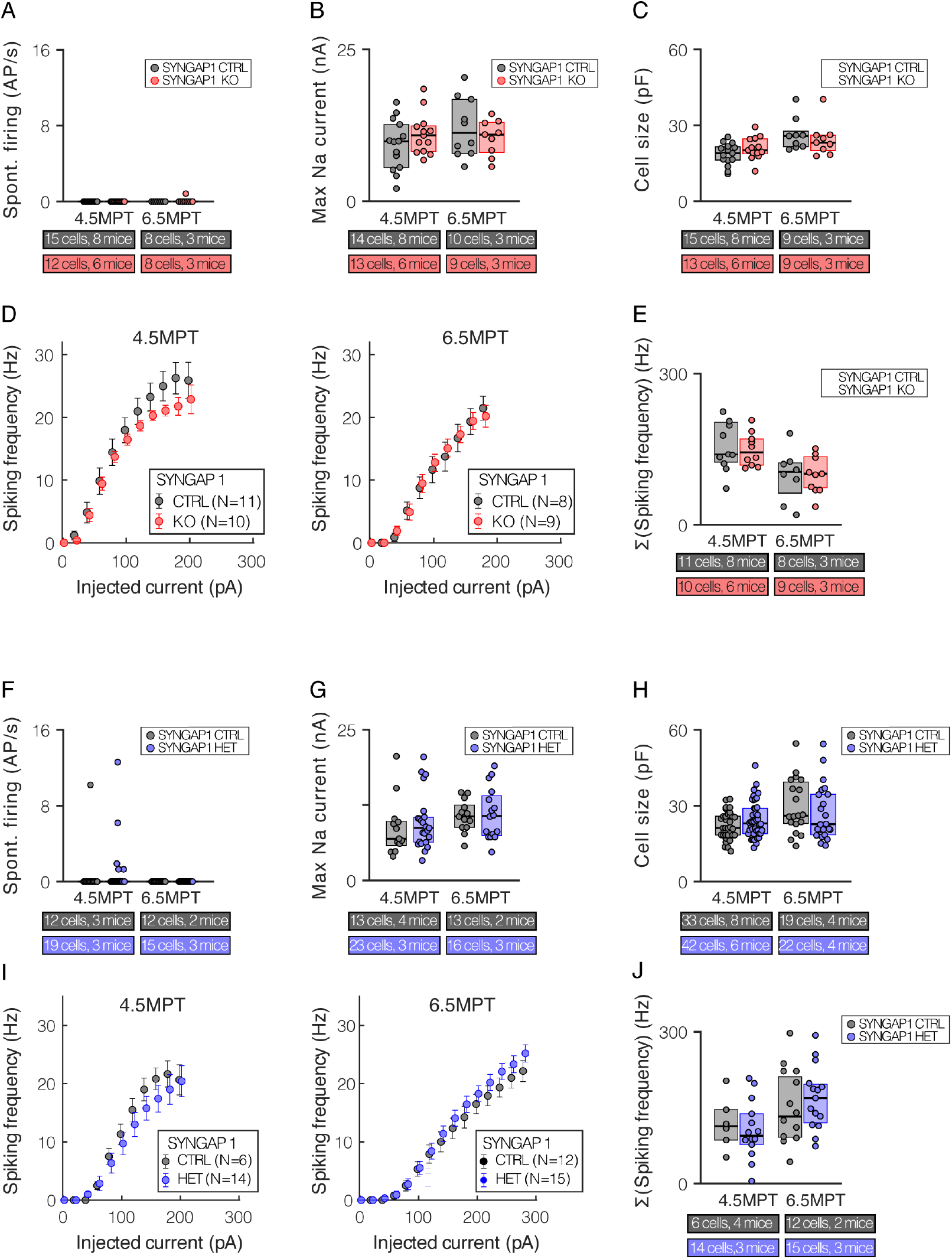
Comparison of intrinsic properties from patch-clamp recordings. **(A-E)** Additional cell intrinsic properties showing no difference between SYNGAP1 CTRL and KO (A) Spontaneous firing rate, (B) Max Sodium current, (C) Cell size, (D) f/I curves and (E) Summed spiking frequency, where we sum all spike from the f/I curve in (D). **(F-J)** Additional cell intrinsic properties showing no difference between SYNGAP1 CTRL and KO (F) Spontaneous firing rate, (G) Max sodium current, (H) Cell size, (I) f/I curves and (J) Summed spiking frequency, as in (E).

**Fig. S3.**
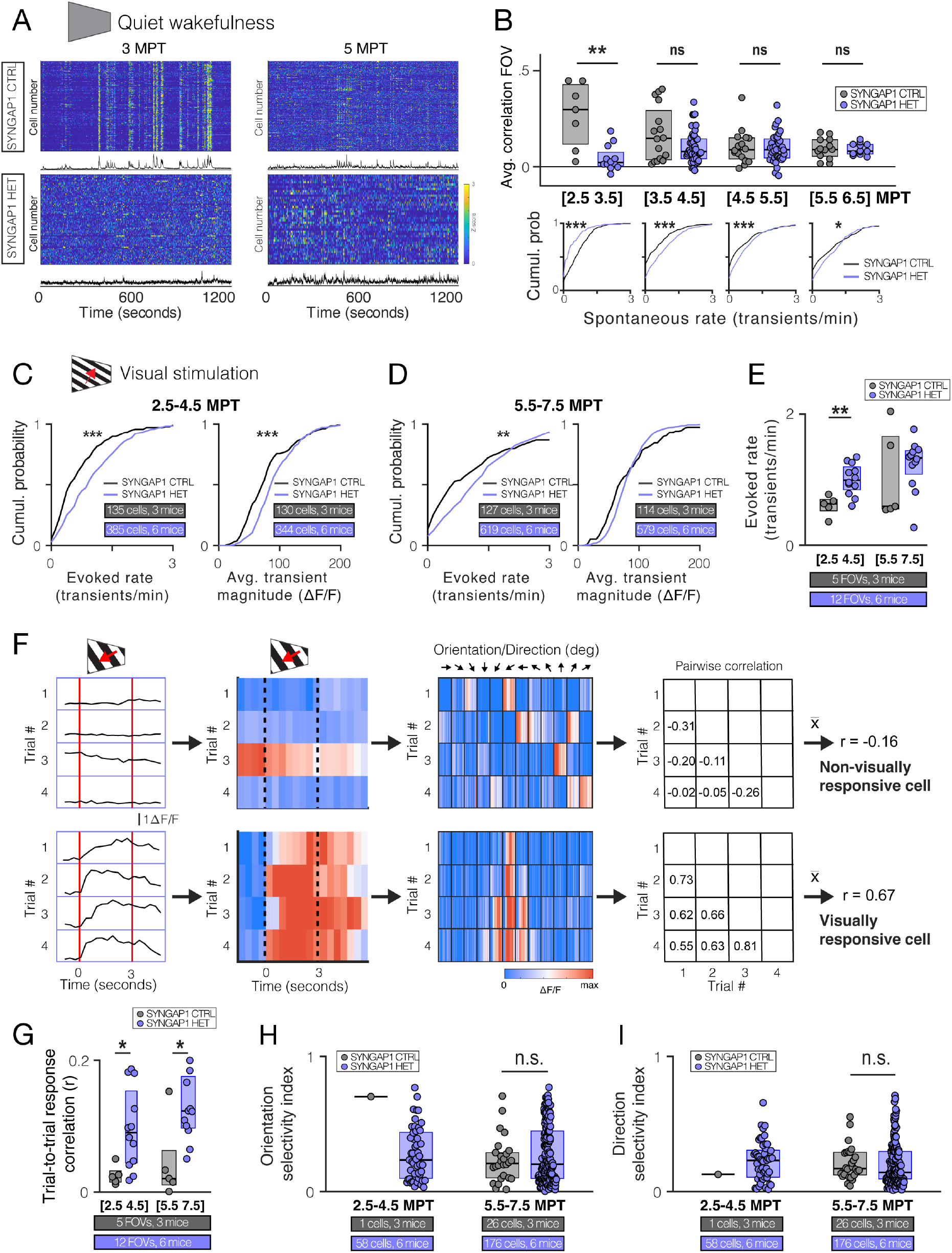
Functional calcium data on spontaneous synchronization and visual tuning. **(A)** Population activity graphs from example FOVs in CTRL and HET at 3 and 5 MPT. Top left shows highly synchronous bursting patterns in CTRL, that disappears over time. HET neurons do not exhibit this type of activity at the times we measured. Values shown are in rows are the Z-scored df/f calcium traces of individual neurons recorded for 20 minutes. No visual stimulus was presented during this time. **(B)** Quantification of correlated activity at 4 different time epochs. *Top:* dots indicate average pairwise correlation between activity traces of all simultaneously recorded neurons for individual FOVs. Note the elevated correlation for CTRL FOVs in the first time epoch. *Bottom:* Cumulative probability plots showing the distribution of rate of calcium transients per neuron for CTRL or HET neurons. Note the reversal of the curves between first and later time epochs. **(C)** Similar to the data shown in main figure 3C,D now for traces recorded during visual stimulation. Here, at the time point 2.5-4.5MPT, we find a strong increase in activity for HET over CTRL, both for transient rate as average transient magnitude. **(D)** Same parameters as quantified in (C) for neurons recorded between 5.5-7.5 MPT. The transient rates for HET neurons is increased compared to CTRL neurons. **(E)** Control analysis where we compare transient rates averaged per FOV for both genotypes across time. This analysis revealed that only at early time points show an increased activity for HET neurons that is consistent per FOVs. **(F)** Outline of the calculation methods for the trial-to-trial response correlation measure we use to quantify the consistency of response to 4 repeated presentations of the same visual stimulus. We extract calcium traces from windows centered on the presentation of each individual visual stimulus (0.5 s before – 0.5 s after stimulus). These are then concatenated per trial for all 12 orientations. The 4 resulting response vectors are correlated in a pair-wise fashion. The average of these correlations (r) is then thresholded to classify a neuron as being visually responsive (threshold = 0.20). **(G)** Control analysis per FOV like shown in (E), this time for trial-to-trial response correlation (r). This analysis revealed that the difference we observe when pooling all neurons is confirmed when averaging values per FOV. **(H)** Summary of orientation tuning index (OSI) and **(I)** direction tuning index (DSI) for neurons passing the threshold defined above. HET neurons are not different from CTRL when we compare these tuning indices, the main difference is the number of neurons that pass the visual tuning threshold (shown in Fig. 3K).

